# Low-cost HPV testing and the prevalence of cervical infection in asymptomatic populations in Guatemala

**DOI:** 10.1101/134775

**Authors:** Hong Lou, Eduardo Gharzouzi, Sarita Polo Guerra, Joël Fokom Domgue, Edmundo Torres-Gonzalez, Julie Sawitzke, Guillermo Villagran, Lisa Garland, Joseph Boland, Sarah Wagner, Joanna Martinez, Héctor Rosas, Jami Troxler, Heidi McMillen, Bailey Kessing, Enrique Alvirez, Miriam Castillo, Hesler Morales, Victor Argueta, Andert Rosingh, Femke J.H.B. van Aerde-van Nunen, Herbert M. Pinedo, Griselda Lopez, Mark Schiffman, Michael Dean, Roberto Orozco

## Abstract

**Background:** A low cost and accurate method for detecting high-risk (HR) HPV is important to permit HPV testing for cervical cancer prevention. We validated a low-cost commercially available HPV method (H13, Hybribio, Hong Kong) and determined the distribution of HPV infections in over 1717 cancer-free women in Guatemala.

**Methods:** H13 results were compared with two more established HPV tests: (Xpert™ (Cepheid) and SPF10-LIPA_25_™ (DDL)) in 40 mainly known positive specimens. HR-HPV was detected in cervical samples from 1717 cancer-free women receiving Pap smears using the Hybribio™ realtime PCR assay of 13 HR types. Selected HPV positive samples were sequenced to determine viral type.

**Results:** The Hybribio H13 Assay showed 93% identical results with Xpert, and 89% with SPF10-LIPA_25_. A total of 13% (226/1717) of women tested HPV+. The highest prevalence was found in younger women (<30 years, 22 %) and older ones (≥60 years, 15%). The six most common HR-HPV types among the 148 HPV+ typed were HPV16 (22%), HPV18 (11%), HPV39 (11%), HPV58 (10%), HPV52 (8%), and HPV45 (8%).

**Conclusions:** The Hybribio Assay was low cost, and accurate in testing for HR-HPV infection. As in most places, HPV16 was the most prevalent HR type in Guatemala and the age-specific prevalence curve peaked in younger ages.

**Highlights:**

1. A low cost and accurate method, Hybribio Assay, could be used for CC testing in low income regions.
2. A total of 13% of cancer-free women were HPV+ and positivity was associated with younger age (<30 years old) in Guatemala.
3. HPV16 was the major prevalent type.

## Introduction

In Guatemala, cervical cancer (CC) is a leading cause of cancer in women (1530 cases/year, Age-standardized rate (ASR) 31/100,000) resulting in an estimated 717 deaths) (Globocan 2012). In the Instituto de Cancerología (INCAN), the main adult oncology hospital in Guatemala, over 40% of women diagnosed with malignancies have advanced CC, requiring costly management that often has a poor outcome (1). Therefore, a focus on prevention is important.

There is overwhelming evidence that persistent infection with specific types of HPV is the main cause of CC (2, 3). HPV types that are classified as established carcinogens are HPV16, 18, 31, 33, 35, 39, 45, 51, 52, 56, 58, 59, and possibly 68 (4). While prophylactic vaccination of adolescents and possibly young adults is the ultimate preventive strategy, screening will remain important for decades to come. Cytology-based screening has been associated with a major reduction in the incidence and mortality of the disease in developed countries (5). However, cytology is either unavailable or poorly conducted in most low-income countries (6).

In most populations, HPV incidence peaks in women in their late teens or early 20s (7-9) following the average age of first sexual intercourse. Incidence and prevalence typically decline from age 30-45 (10). A second peak is often observed in postmenopausal women, possibly related to immune senescence and escape from long-term cell-mediated immune control of infections acquired earlier in life (11, 12). HPV DNA testing has been proposed as an alternative to cytology in women older than 25-30 years, when prevalence declines and predictive value (for signaling risk of precancer) of a positive HPV test increases. (13-15). However, commercially available tests are typically expensive and require sophisticated equipment (16, 17). The use of HPV assays targeting lower-resource settings would be useful for CC prevention in such settings, which contribute most of the world CC burden.

This study sought to assess the concordance of a new low-cost HPV assay (H13, Hybribio Ltd, Hong Kong) with more established commercial tests, GeneXpert (Xpert, Cepheid, California, USA) and SPF10-LIPA_25_ (DDL, Netherlands) assays (18-20). In addition, we determined the prevalence of HPV infection in women attending screening clinics in Guatemala.

## Materials and Methods

### Study populations

To validate the H13 test against other clinical tests, a convenience sample of 40 mostly known HPV positive subjects who had undergone cervical screening and testing at the Curacao Medical Lab was employed. To explore the epidemiology of HPV in Guatemala using H13, a total of 1717 samples were obtained from asymptomatic sexually active women undergoing routine screening with Pap smear at hospital-based screening clinics in Guatemala, after obtaining informed consent. A questionnaire on reproductive and socio-demographic characteristics was administered by trained personnel on the Guatemalan subjects (21). The samples were collected by a medical practitioner using a Dacron swab placed in a tube containing 3 ml of Scope mouthwash, maintained at 4°C and transported at room temperature (22). The study was approved by an Institutional Review Board (IRB) and testing in the US laboratory was judged exempt by the NIH Office of Human Studies Research.

### HPV testing

The detection of HPV was performed using a Hybribio Assay that detects the13 high-risk HPV types by Real time PCR (Hybribio Biochem Co. Ltd. China). A cell lysate was prepared per the manufacturer’s instruction. To determine HPV type, selected HPV positive samples were subjected to Touchdown PCR and DNA sequencing (Fig. 2). Samples in which the internal control (IC) did not amplify on two separate attempts were excluded (171 samples).

The sensitivity of the Touchdown PCR was determined by a series of 10-fold dilutions of DNA from HPV+ and HPV-cell lines using Broad-Spectrum (BS) GP5+ and GP6+ Primers (BSGP5+/6+) (150bp) (Supplementary Fig. 1) (23). For the Hybribio Assay, each 96-well plate included four HPV+ controls: CC cell line DNA from HeLa (HPV18), CaSki (HPV16), MS751 (HPV45) DNA and positive control DNA from the Hybribio Assay kit; HPV-controls included C33A DNA and water. The HPV+ samples were amplified by using 400 nM BSGP5+/6+. Briefly, 10 min denaturation step at 95°C was followed by 40 cycles of amplification. Each cycle included denaturation at 94°C for 20 s, annealing at 38°C for 30 s, and elongation at 71° C for 80 s. The final elongation step was 5 min. The ramping rates were adjusted as described (24); 1.8° C/cycle from 74° C to 38° C in first 20 cycles. Each experiment included HPV+ and HPV-controls and a sample lacking template DNA (Supplementary Fig. 2). The PCR products were subjected to Sanger Sequencing on an ABI3730XL. Sequences were analyzed by assembly and trimming in SeqMan (DNASTAR, Madison, WI) followed by BLAST search (NCBI). Samples with inconclusive Sanger sequence were repeated with a next-generation sequencing method (S. Wagner, and J. Boland, manuscript in preparation).

### Statistical analyses

We compared the detection of HPV by the Hybribio Assay to Xpert and SPF10-LIPA_25_ in 40 cervical cytology samples collected in a clinical laboratory in Curacao and calculated percent agreement and kappa. Statistical analyses on the Guatemalan samples were performed to determine age-specific HPV prevalence, comparing the age groups with the Pearson Chi-square test using GraphPad Prism version 7 for Windows. P < 0.05 was regarded as statistically significant. We performed analyses of association between HPV infection and age.

## Results

To evaluate the Hybribio H13 test as a potential low cost assay for HR-HPV testing we compared it against two commercial assays, Xpert and DDL. Table 1 show that nearly identical results were seen between Hybribio Assay and the other tests (93%). Also, 89% identical results were observed between SPF10-LIPA_25_ and Xpert (**Table 1**). We also evaluated the sensitivity and required assay volume of the H13 test and determined that a 10ul real-time PCR volume gave equivalent results (**Table 2**). In a separate study, we determined that the H13 test give high concordance with the Qiagen HC2 and Onclarity assays, and has good clinical accuracy compared to histologic diagnosis (25). Therefore, H13 is a low-cost and reliable assay for HPV screening in settings where aggregate detection of all 13 high-risk types is desired.

**Table 1.** Comparison of the Hybribio Assay to the Xpert and SPF10-LIPA_25_ assay

|  | HR HPV, No. |  |  |  | Overall Agreement (%) | Kappa |
| --- | --- | --- | --- | --- | --- | --- |
|  | Pos/Pos | Pos/Neg | Neg/Pos | Neg/Neg |  |  |
| HybriBio Assay/Xpert | 30 | 2 | 1 | 7 | 93% | 0.78 |
| HybriBio Assay/SPF10-LIPA<sub>25</sub>* | 22 | 0 | 2 | 4 | 93% | 0.76 |
| SPF10-LIPA<sub>25</sub>/Xpert | 21 | 3 | 0 | 4 | 89% | 0.67 |

Data shown for the 13 HR types in common to the three assays. HR, high risk; HPV, human papillomavirus; Pos, positive; Neg, negative.

**Table 2.** Comparison of the Hybribio Assay with 10 μl to 20 μl volume

|  |  |  |  | 13HR HPV Types<br>(FAM) | Internal Control<br>(JOE) | HPV<br>Status |
| --- | --- | --- | --- | --- | --- | --- |
| Sample | Total Volume (μl) | Input DNA (5ng/μl)<br>Volume (μl) | Input DNA<br>(ng) | Ct | Ct |  |
| HPV positive controls |  |  |  |  |  |  |
| CaSki (HPV16) | 20 | 2 | 10 | 15.74 | 27.26 | + |
|  | 10 | 1 | 5 | 16.15 | 28.61 | + |
| HeLa (HPV18) | 20 | 2 | 10 | 22.26 | 26.23 | + |
|  | 10 | 1 | 5 | 23.03 | 26.26 | + |
| MS751 (HPV45) | 20 | 2 | 10 | 26.90 | 29.79 | + |
|  | 10 | 1 | 5 | 27.96 | 29.91 | + |
| ME180 (HPV39/18) | 20 | 2 | 10 | 27.61 | 25.99 | + |
|  | 10 | 1 | 5 | 27.72 | 25.96 | + |
| Unknown samples from cell lysate |  |  |  |  |  |  |
| S1 DD015959 | 20 | 2 |  | ND* | 24.05 | - |
|  | 10 | 1 |  | ND | 24.80 | - |
| S2 DD015960 | 20 | 2 |  | 17.60 | 27.59 | + |
|  | 10 | 1 |  | 18.02 | 28.10 | + |
| S3 DD015963 | 20 | 2 |  | 16.85 | 20.78 | + |
|  | 10 | 1 |  | 17.20 | 21.30 | + |
| S6 DD015964 | 20 | 2 |  | 16.01 | 22.05 | + |
|  | 10 | 1 |  | 16.80 | 22.70 | + |
| HPV negative control (C33A) | 20 | 2 | 10 | ND | 25.66 | - |
|  | 10 | 1 | 5 | ND | 25.04 | - |
| Controls from kit |  |  |  |  |  |  |
| Positive control (Kit) | 20 | 2 |  | 23.88 | 23.39 | + |
|  | 10 | 1 |  | 25.07 | 25.68 | + |
| Negative control (Kit)** | 20 | 2 |  | ND | ND | - |
|  | 10 | 1 |  | ND | ND | - |

\*: Not detected
\*\*: DNase-free distilled water from kit

To apply the H13 test to samples from a low-middle income country, we used the test to determine the prevalence and distribution of HPV types in Guatemala. We recruited asymptomatic women from the general population at hospital clinics performing cytology screening. The women sampled ranged in age from 17 to 79 years attending clinics in Guatemala City and the city of Puerto Barrios. To determine the prevalence of HPV infection, 1717 subjects were tested (Fig. 2). The overall prevalence of HPV infection was 13% (226/1717) (Table 4). To understand the age specific prevalence of HPV infection, the women were divided into 6 age groups (Table 3). HPV prevalence ranged from 22% in the <30 group to 8% to 14% in the 30-59 age groups and 15% in the ≥60 group. Age specific prevalence for HPV was significantly higher in the younger age groups (<30) (P=0.0022) (Table 3 and Figure 1).

**Figure 1.**
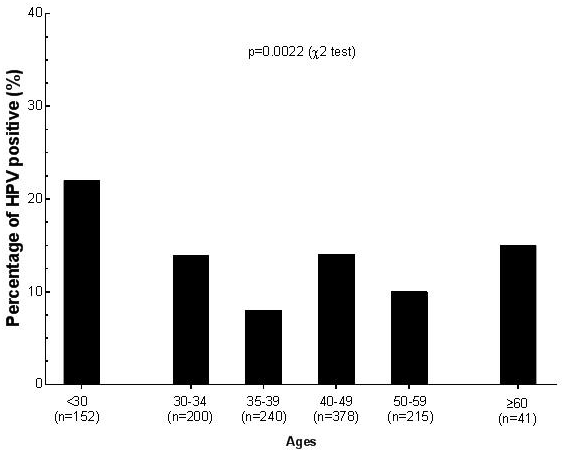
Age specific prevalence of HPV by age group in Guatemala. The prevalence of HPV in asymptomatic women in Guatemala is displayed by age group. A Pearson test of all six groups is statistically significant (P<0.0022). The number of total women in each age group (n) is shown.

**Figure 2.**
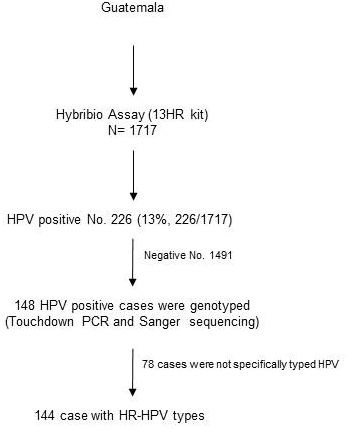
Flow chart of participants and associated HPV test outcomes.

**Table 3.** Analysis of age risk factors related to 13HR HPV detection by Hybribio assay

| Age group | Total No. | HPV+ No. | HPV+ (%) | OR | 95% CI | *P value* |
| --- | --- | --- | --- | --- | --- | --- |
| <30 | 152 | 34 | 22% |  |  |  |
| 30-34 | 200 | 27 | 14% | 0.542 | 0.3147 to 0.9492 | 0.0294 |
| 35-39 | 240 | 20 | 8% | 0.316 | 0.1732 to 0.5627 | 0.0001 |
| 40-49 | 378 | 52 | 14% | 0.554 | 0.3458 to 0.8866 | 0.015 |
| 50-59 | 215 | 21 | 10% | 0.376 | 0.2099 to 0.6647 | 0.0009 |
| ≥60 | 41 | 6 | 15% | 0.595 | 0.2400 to 1.5410 | 0.2782 |

**Table 4.**
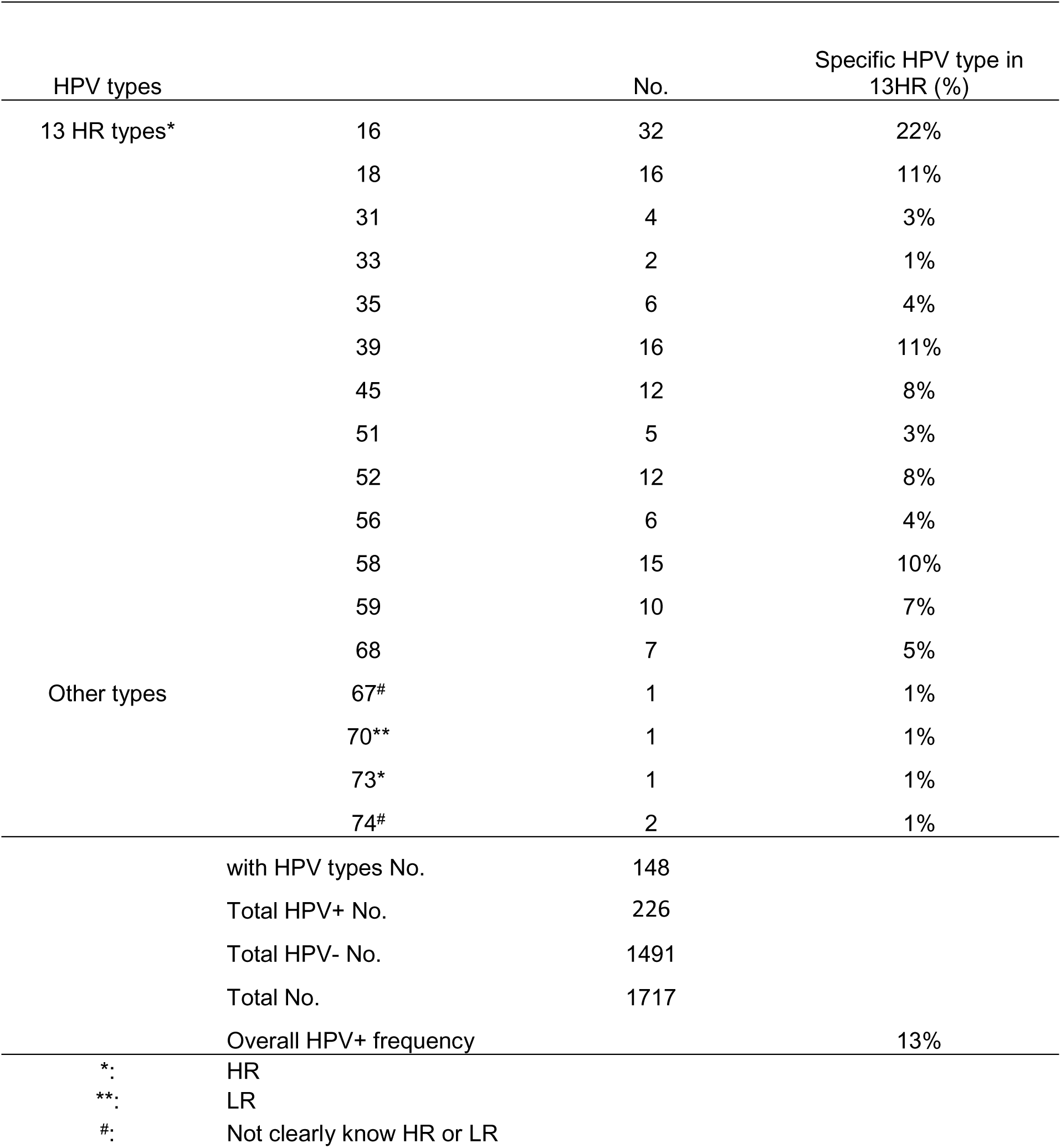
Prevalence of HR-HPV types by Hybribio assay in cancer free women in Guatemala

HPV typing was successful in 148 HPV+ samples and the 13 HR-HPV types 16, 18, 31, 33, 35, 39, 45, 51, 52, 56, 58, 59, 68 were detected in 143 subjects (Table 3.) (143/226). Of the 13 HR types, HPV16 (22%, 32/148), HPV18 (11%, 16/148) accounted for 32% (48/148) and HPV39, 58, 45, 52 combined accounted for 37%, 55/148). Types considered to convey very low risk (Unk), HPV67 and 74 were detected in 1% (Table 4).

## Discussion

Numerous studies support HPV testing as the most sensitive primary screening method for CC. However, there are few HPV tests that are affordable for LMICs. We performed a small comparison study of the Hybribio H13 test with two other commercial assays, GeneXpert and DDL spf10LiPA and demonstrated a high rate of concordance. Commercial tests in Guatemala cost between $100-210, out of the range of practical use. Mexico carried out a large screening program with the hybrid capture assay (HC2, Qiagen, Germantown, MD) at an approximate cost of $11 per test, but many poor and rural areas remain unscreened. A method has been developed with support from donor and non-profit foundations, called careHPV (Qiagen). This system has been rigorously tested in China, India and in pilot programs in other areas (26) and is under further evaluation in several Latin American countries. It is unclear whether it can be scaled up to cover a whole population. Thus, there is still considerable discussion on the most effective strategies for managing HPV+ women in different economic and cultural settings (27-29).

We sought to establish a method that would be cost effective, and use only equipment available in a standard molecular biology laboratory. We employed a validated storage buffer, mouthwash containing 15% ethanol, which costs $0.01 per sample with fewer shipping requirements than methanol based buffers (22). The Hybribio Assay under the conditions we employed (10ul reaction volume), costs about $3/assay (Table 2). While Hybribio requires a real-time PCR instrument, we have purchased used ABI7000 instruments for under $1000, and found that some Latin American hospitals and clinical laboratories have real-time PCR instrumentation.

In this study, the Hybribio Assay was compared for detection of HPV to two clinically validated tests (Xpert and SPF10-LIPA_25_). Both Hybribio H13 and Xpert target the E6 and E7 of 13 and 14 HR-HPV types, respectively. Hybribio Assay has a pooled probe for HR-HPV types, 16, 18, 31, 33, 35, 39, 45, 51, 52, 56, 58, 59, 68; whereas Xpert test detects HR-HPV16, 18/45 in separate detection channels, with 11 other HR types detected in 3 additional channels. HR types detected are the same as the Hybribio Assay except HPV66 is included [19, 20]. The SPF10-LIPA_25_ test targets the conserved L1 region and detects 14 HR types and 11 LR types (18). In a separate study, we further validated H13 against HC2 (Qiagen) and Onclarity (BD Diagnostics, Sparks, MD) ((25)).

Few studies of HPV prevalence have been described in Guatemala, even though this country has one of the highest incidences of HPV related diseases and mortality in Latin America (1, 30). However, our study has the advantage of collecting samples from healthy women using a low-cost HPV screening method. The overall prevalence of HR HPV infection in this study was 13% for Guatemala, similar to rural Costa Rica (31) and lower than studies from Northern Spain (29%) (7) and other countries (25% to 29%) (32, 33). The difference is likely due in part to the greater age (thus, lower prevalence) of our study population.

As seen in most countries, HPV16 is the most common type found in women with or without cancer (34, 35). The combined prevalence of HPV16, and HPV18 was highest in the youngest age groups in this country. A similar prevalence of HPV16 and 18 was reported in other studies (36) as well as in Guatemalans with CC (21). In addition to HPV16 and 18, 15 other HPV types were observed frequently in our study, most notably HR-HPV 39, 58, 45, and 52. The relatively small number of women <30 years of age had the highest prevalence, while the HPV prevalence decreased markedly with increasing age, up to age 60. This trend has been observed in Costa Rica and many other studies (7-9).

Our study has several limitations that might affect our conclusions. We used a Hybribio Assay that detects only 13 HR types. In addition, we attempted to sequence all positive samples to determine type. In a small portion of samples, we had a failure to detect the IC indicating a failure in sample collection, preservation or storage. Most of these samples were negative using both Hybribio and a touchdown PCR method (data not shown). This could indicate a limitation to using Scope mouthwash as a preservative. We have limited cytology data on the women and have not demonstrated that Hybribio Assay is effective in CC prevention in Guatemala. However, the manufacturer reports data on a comparison with Qiagen (with >95% agreement with an FDA approved test), the test passed two WHO proficiency trials and has been used in a study of 48,559 women in China (11).

In conclusion, we have established the Hybribio H13 test as an affordable alternative for HPV screening. HPV infection was detected in 13% of asymptomatic women in Guatemala. The distribution of HPV types is typical of other countries and the highest HPV prevalence is in youngest age groups (<30). This low-cost approach to detect HPV could be employed in other countries planning to introduce HPV screening to reduce the burden of CC.

## Acknowledgements

The authors would like to thank the staff and health professionals from the Instituto de Cancerologia, Guatemala City, Guatemala, Hospital San Juan de Dios, Guatemala City, Guatemala and Medical Laboratory Services, Willemstad, Curacao, as well as Funashon Prevenshon, Curacao, and the Liga Nacional Contra Cancer, Guatemala. Patricia Zaid, Adolfo Santizo, Esther Avila and Lineth Boror for sample and data collection and shipping; Russ Hanson for approvals, Qi Yang for statistical advice, Bob Pinedo and Vicky de Falla for helpful discussions and the BSP-CCR Genetics Core for technical support. Supported in part by the Intramural Research Program of the National Institutes of Health, National Cancer Institute, Center for Cancer Research, and Division of Cancer Epidemiology and Genetics from Leidos-Frederick under contract # HHSN261200800001E. The content of this publication does not necessarily reflect the views or policies of the Department of Health and Human Services, nor does mention of trade names, commercial products, or organizations imply endorsement by the U.S. government.

## References

1. Valles X, Murga GB, Hernandez G, Sabido M, Chuy A, Lloveras B, Alameda F, de San Jose S, Bosch FX, Pedroza I, Castellsague X, Casabona J. 2009. High prevalence of human papillomavirus infection in the female population of Guatemala. Int J Cancer 125:1161-7.

2. Ngelangel C, Munoz N, Bosch FX, Limson GM, Festin MR, Deacon J, Jacobs MV, Santamaria M, Meijer CJ, Walboomers JM. 1998. Causes of cervical cancer in the Philippines: a case-control study. J Natl Cancer Inst 90:43-9.

3. Smith JS, Lindsay L, Hoots B, Keys J, Franceschi S, Winer R, Clifford GM. 2007. Human papillomavirus type distribution in invasive cervical cancer and high-grade cervical lesions: a meta-analysis update. Int J Cancer 121:621-32.

4. Bouvard V, Baan R, Straif K, Grosse Y, Secretan B, El Ghissassi F, Benbrahim-Tallaa L, Guha N, Freeman C, Galichet L, Cogliano V. 2009. A review of human carcinogens–Part B: biological agents. Lancet Oncol 10:321-2.

5. Arbyn M, Raifu AO, Weiderpass E, Bray F, Anttila A. 2009. Trends of cervical cancer mortality in the member states of the European Union. Eur J Cancer 45:2640-8.

6. Gakidou E, Nordhagen S, Obermeyer Z. 2008. Coverage of cervical cancer screening in 57 countries: low average levels and large inequalities. PLoS Med 5:e132.

7. de Ona M, Alvarez-Arguelles ME, Torrents M, Villa L, Rodriguez-Feijoo A, Palacio A, Boga JA, Tamargo A, Melon S. 2010. Prevalence, evolution, and features of infection with human papillomavirus: a 15-year longitudinal study of routine screening of a women population in the north of Spain. J Med Virol 82:597-604.

8. Leinonen MK, Anttila A, Malila N, Dillner J, Forslund O, Nieminen P. 2013. Type- and age-specific distribution of human papillomavirus in women attending cervical cancer screening in Finland. Br J Cancer 109:2941-50.

9. Ley C, Bauer HM, Reingold A, Schiffman MH, Chambers JC, Tashiro CJ, Manos MM. 1991. Determinants of genital human papillomavirus infection in young women. J Natl Cancer Inst 83:997-1003.

10. Schiffman M, Doorbar J, Wentzensen N, de Sanjose S, Fakhry C, Monk BJ, Stanley MA, Franceschi S. 2016. Carcinogenic human papillomavirus infection. Nat Rev Dis Primers 2:16086.

11. Chen Q, Xie LX, Qing ZR, Li LJ, Luo ZY, Lin M, Zhang SM, Chen WZ, Lin BZ, Lin QL, Li H, Chen WP, Zheng PY, Mao LZ, Chen CY, Yang C, Zhan YZ, Liu XZ, Zheng JK, Yang LY. 2012. Epidemiologic characterization of human papillomavirus infection in rural Chaozhou, eastern Guangdong Province of China. PLoS One 7:e32149.

12. Syrjanen K. 2008. New concepts on risk factors of HPV and novel screening strategies for cervical cancer precursors. Eur J Gynaecol Oncol 29:205-21.

13. Mayrand MH, Duarte-Franco E, Rodrigues I, Walter SD, Hanley J, Ferenczy A, Ratnam S, Coutlee F, Franco EL. 2007. Human papillomavirus DNA versus Papanicolaou screening tests for cervical cancer. N Engl J Med 357:1579-88.

14. Naucler P, Ryd W, Tornberg S, Strand A, Wadell G, Elfgren K, Radberg T, Strander B, Johansson B, Forslund O, Hansson BG, Rylander E, Dillner J. 2007. Human papillomavirus and Papanicolaou tests to screen for cervical cancer. N Engl J Med 357:1589-97.

15. Rijkaart DC, Berkhof J, Rozendaal L, van Kemenade FJ, Bulkmans NW, Heideman DA, Kenter GG, Cuzick J, Snijders PJ, Meijer CJ. 2012. Human papillomavirus testing for the detection of high-grade cervical intraepithelial neoplasia and cancer: final results of the POBASCAM randomised controlled trial. Lancet Oncol 13:78-88.

16. Barski A, Cuddapah S, Cui K, Roh TY, Schones DE, Wang Z, Wei G, Chepelev I, Zhao K. 2007. High-resolution profiling of histone methylations in the human genome. Cell 129:823-37.

17. Green RE, Krause J, Ptak SE, Briggs AW, Ronan MT, Simons JF, Du L, Egholm M, Rothberg JM, Paunovic M, Paabo S. 2006. Analysis of one million base pairs of Neanderthal DNA. Nature 444:330-6.

18. Castle PE, Porras C, Quint WG, Rodriguez AC, Schiffman M, Gravitt PE, Gonzalez P, Katki HA, Silva S, Freer E, Van Doorn LJ, Jimenez S, Herrero R, Hildesheim A. 2008. Comparison of two PCR-based human papillomavirus genotyping methods. J Clin Microbiol 46:3437-45.

19. Cuschieri K, Geraets D, Cuzick J, Cadman L, Moore C, Vanden Broeck D, Padalko E, Quint W, Arbyn M. 2016. Performance of a Cartridge-Based Assay for Detection of Clinically Significant Human Papillomavirus (HPV) Infection: Lessons from VALGENT (Validation of HPV Genotyping Tests). J Clin Microbiol 54:2337-42.

20. Einstein MH, Smith KM, Davis TE, Schmeler KM, Ferris DG, Savage AH, Gray JE, Stoler MH, Wright TC, Jr., Ferenczy A, Castle PE. 2014. Clinical evaluation of the cartridge-based GeneXpert human papillomavirus assay in women referred for colposcopy. J Clin Microbiol 52:2089-95.

21. Lou H, Villagran G, Boland JF, Im KM, Polo S, Zhou W, Odey U, Juarez-Torres E, Medina-Martinez I, Roman-Basaure E, Mitchell J, Roberson D, Sawitzke J, Garland L, Rodriguez-Herrera M, Wells D, Troyer J, Pinto FC, Bass S, Zhang X, Castillo M, Gold B, Morales H, Yeager M, Berumen J, Alvirez E, Gharzouzi E, Dean M. 2015. Genome Analysis of Latin American Cervical Cancer: Frequent Activation of the PIK3CA Pathway. Clin Cancer Res 21:5360-70.

22. Castle PE, Sadorra M, Garcia FA, Cullen AP, Lorincz AT, Mitchell AL, Whitby D, Chuke R, Kornegay JR. 2007. Mouthwash as a low-cost and safe specimen transport medium for human papillomavirus DNA testing of cervicovaginal specimens. Cancer Epidemiol Biomarkers Prev 16:840-3.

23. Schmitt M, Dondog B, Waterboer T, Pawlita M. 2008. Homogeneous amplification of genital human alpha papillomaviruses by PCR using novel broad-spectrum GP5+ and GP6+ primers. J Clin Microbiol 46:1050-9.

24. Snijders PJ, van den Brule AJ, Jacobs MV, Pol RP, Meijer CJ. 2005. HPV DNA detection and typing in cervical scrapes. Methods Mol Med 119:101-14.

25. Fokom Domgue J, Schiffman M, Wentzensen NH, Gage JC, Castle PE, Raine-Bennett TR, Fetterman B, Lorey T, Poitras NE, Befano B, Xie Y, Miachon LS, Dean M. 2017. Assessment of a new lower-cost real-time PCR detection assay for high-risk HPV: Useful for cervical screening in limited resource settings? J Clin Microbiol doi:10.1128/JCM.00492-17.

26. Trope LA, Chumworathayi B, Blumenthal PD. 2009. Preventing cervical cancer: stakeholder attitudes toward CareHPV-focused screening programs in Roi-et Province, Thailand. Int J Gynecol Cancer 19:1432-8.

27. Gottschlich A, Rivera-Andrade A, Grajeda E, Alvarez C, Mendoza Montano, Meza R. 2017. Acceptability of Human Papillomavirus Self-Sampling for Cervical Cancer Screening in an Indigenous Community in Guatemala. Journal of Global Oncology 0:JGO.2016.005629.

28. Jeronimo J, Holme F, Slavkovsky R, Camel C. 2016. Implementation of HPV testing in Latin America. J Clin Virol 76 Suppl 1:S69-73.

29. Schiffman M, Wentzensen N, Wacholder S, Kinney W, Gage JC, Castle PE. 2011. Human papillomavirus testing in the prevention of cervical cancer. J Natl Cancer Inst 103:368-83.

30. Nunez-Troconis J, Delgado M, Gonzalez J, Mindiola R, Velasquez J, Conde B, Whitby D, Munroe DJ. 2009. Prevalence and risk factors of human papillomavirus infection in asymptomatic women in a Venezuelan urban area. Invest Clin 50:203-12.

31. Herrero R, Hildesheim A, Bratti C, Sherman ME, Hutchinson M, Morales J, Balmaceda I, Greenberg MD, Alfaro M, Burk RD, Wacholder S, Plummer M, Schiffman M. 2000. Population-based study of human papillomavirus infection and cervical neoplasia in rural Costa Rica. J Natl Cancer Inst 92:464-74.

32. de Sanjose S, Diaz M, Castellsague X, Clifford G, Bruni L, Munoz N, Bosch FX. 2007. Worldwide prevalence and genotype distribution of cervical human papillomavirus DNA in women with normal cytology: a meta-analysis. Lancet Infect Dis 7:453-9.

33. Dunne EF, Unger ER, Sternberg M, McQuillan G, Swan DC, Patel SS, Markowitz LE. 2007. Prevalence of HPV infection among females in the United States. Jama 297:813-9.

34. Alibegashvili T, Clifford GM, Vaccarella S, Baidoshvili A, Gogiashvili L, Tsagareli Z, Kureli I, Snijders PJ, Heideman DA, van Kemenade FJ, Meijer CJ, Kordzaia D, Franceschi S. 2011. Human papillomavirus infection in women with and without cervical cancer in Tbilisi, Georgia. Cancer Epidemiol 35:465-70.

35. Zhao R, Zhang WY, Wu MH, Zhang SW, Pan J, Zhu L, Zhang YP, Li H, Gu YS, Liu XZ. 2009. Human papillomavirus infection in Beijing, People’s Republic of China: a population-based study. Br J Cancer 101:1635-40.

36. Haguenoer K, Giraudeau B, Gaudy-Graffin C, de Pinieux I, Dubois F, Trignol-Viguier N, Viguier J, Marret H, Goudeau A. 2014. Accuracy of dry vaginal self-sampling for detecting high-risk human papillomavirus infection in cervical cancer screening: a cross-sectional study. Gynecol Oncol 134:302-8.

